# Density dependence shapes life-history trade-offs in a food limited ungulate population

**DOI:** 10.1101/2024.01.26.577325

**Authors:** Harman Jaggi, Wenyun Zuo, Rosemarie Kentie, Jean-Michel Gaillard, Tim Coulson, Shripad Tuljapurkar

## Abstract

Quantifying trade-offs within populations is an important goal in life-history and evolutionary theory. However, most studies focusing on life-history variation assume trade-offs to be static. In this paper, we provide a framework for understanding life-history variation at different densities while enabling us to reveal trade-offs that are often masked due to individual heterogeneity. We use published individual-based data from a population of Soay sheep and find density dependence strongly shapes life-history trade-offs and the distribution of lifetime reproductive success. We find trade-offs between juvenile survival and growth structures life-history variation for Soay sheep at all densities, but the intensity of the trade-offs increases with population density. At carrying capacity (*K*), a new trade-off appears between reproduction and juvenile survival. In addition, we find the distribution of LRS to be highly constrained at *K*, with mothers of prime adult sizes (∼ 25kg) contributing the most to reproduction. We also find the effects of density-dependence on demographic measures such as net reproductive rate, stable size distribution, and average vital rate functions. Our results suggest high density limits the diversity of individual life history strategies and has implications for better understanding the evolution of reproductive tactics via density-dependent selection and management of ungulates in food limited environments.

## 1 Introduction

There is tremendous variation in life-history strategies (Stearns 1992; Roff 1993). The evolution of these strategies is known to be constrained by physiological limitations and in food limited populations by trade-offs in resource allocation (Stearns 1989; Descamps et al. 2016). Evolutionary theory posits that organisms must allocate limited resources towards different suite of traits to optimize their fitness thereby resulting in trade-offs. Despite evidence for trade-offs, e.g., offspring-size/number (Smith and Fretwell 1974; Venable 1992), immunocompetence and life-history traits in birds (Norris and Evans 2000), trade-offs observed in natural systems are less ubiquitous than those hypothesized by life-history theory (Metcalf 2016). Trade-offs may be difficult to detect without experimental manipulations because individual variation in resource acquisition is often larger than variation in resource allocation (Van Noordwijk and De Jong 1986), which often masks life-history constraints in empirical studies. Life-history theory predicts costs of reproduction and therefore negative correlations among certain traits. However, individuals can show positive, negative or zero correlations (due to individual heterogeneity or environmental stochasticity) while simultaneously being involved in a classical physiological trade-off (Bell and Koufopanou 1991; Horvitz et al. 1997; Zera and Harshman 2001; Hodgson and Townley 2004; Descamps et al. 2016).

Whether trade-offs within a population are static or change with environmental or demographic conditions is an important question. For instance, as population density varies, individual fitness (ability to survive and reproduce) may also vary thereby impacting life-history outcomes. This could result from chance encounters with potential mates or competitive individuals. Thus, variation in density effects can generate time-varying selection for different life-history strategies (via density-dependent selection). In their recent review, Travis et al. (2023) emphasize the important role of density-dependent selection in evolutionary biology and ecology. The concept that selection on life-history traits could vary with population density has been discussed both empirically and in theory Dobzhansky (1950), MacArthur (1962), MacArthur and Wilson (2001), and Roughgarden (1971).

Pianka (1970) proposed the *r* − *K* selection continuum, where *r* (intrinsic population growth rate) corresponds to no density effects and strategies that optimize putting all energy into reproduction, and producing more offspring; *K* (equilibrium carrying capacity) represents maximal density effects, and strategies that channel energy into maintenance and production of a few fit offspring.

This brings us to our central question-how do we understand life-history variation at different densities while identifying life-history trade-offs (often masked due to individual variation in resource acquisition). Although we often tend to think of trade-offs and density effects separately, we show here that they are intimately linked. We examine trade-offs in a food limited ungulate population by considering the long-term observational data for Soay sheep (*Ovis aries*) in the St. Kilda archipelago, Scotland. The sheep system is of interest because their population has been shown to fluctuate dramatically thereby exhibiting potential for density-dependent selection. Coulson et al. (2001) noted population declines of up to 60% occurred when population size was large and winter weather was harsh. This is due to nonlinear interactions between winter weather, population’s response to density dependence, and food availability (Grenfell et al. 1992; Clutton-Brock et al. 1996; Coulson et al. 1999). The life-history and physiological mechanisms operating at low-density may differ from those operating at high-densities (under intense resource limitation), altering the nature and direction of trade-offs. For instance, at low densities Trinidad guppies allocate more resources to juvenile survival than costs of reproduction (Reznick et al. 2002).

Our first result is that density dependence shapes life-history trade-offs in a food limited ungulate population. We find strong correlations between juvenile survival and growth increment (defined in the methods section). However, the intensity of the negative association between juvenile survival and growth steadily increases with increasing density In addition, a new trade-off shows up between reproduction and juvenile survival with increase in density. Trade-offs are revealed by performing Principal Component Analysis (PCA) on average vital rate functions (survival, recruitment, growth increment). We find changes in population density affect the relationships between vital rates such as, survival, growth and fecundity. The strength of density-dependent responses of vital rates varies a lot, thereby resulting in trade-offs. Our findings are consistent with life-history theory (Stearns 1992) which predicts trade-offs prevent individuals from being proficient at both surviving and growing to large sizes. In addition, Kentie et al. (2020) found Soay sheep individuals with an optimal life history strategy at high-density were different to those having an optimal strategy at low-density and thus individuals could not maximise fitness for both high- and low-density environments.

Next, we examined the effects of density on variation in demographic measures. The demographic measures of interest are- variation in average vital rate functions, net reproductive rate *R*_0_, and stable age-stage distribution (SSD). We also calculated the distribution of lifetime reproductive success (LRS) using Tuljapurkar et al. (2020) to tease apart trade-offs between allocation in survival and costs paid in future reproduction. Many empirical studies have consistently revealed that distribution of LRS are often non-normal, zero-inflated and highly skewed (Cabana and Kramer 1991; Tatarenkov et al. 2008). This begs the question does the distribution of LRS and the skew remain consistent across all population density regimes?

We find the distribution of LRS to be highly constrained at high-densities. LRS is an important component of individual fitness and our research predicts density dependence strongly affects the distribution of LRS. In particular, it is the females at prime adult size (∼ 25kg) that make the most contribution at high densities whereas at low densities contributions come from a wider range of body size (∼ 14-25kg). We find a similar pattern for average vital rate functions and stable stage distribution. We base our model and results on previously published Integral projection model (IPM) and delifing method presented in Coulson (2012) and Coulson et al. (2006), respectively. Using empirical data and life-history trade-offs between IPM parameters, Kentie et al. (2020) created a covariance matrix that allows us to generate strategies spanning the range of possible individual life-histories that Soay sheep are expected to follow.

The work allows us to examine trade-offs, variation in LRS at low and high densities, and comment on how density can operate. Characterizing trade-offs as a function of population density can not only help elucidate the various determinants of observed life-history variation but will also determine what strategies and management implementations may be most relevant for their persistence of the species at a given time. We now describe a brief outline for the paper. The next section introduces the model and methods, followed by our results. We close with a discussion of implications of the research as it provides a framework to study the interplay between density-dependence, lifehistory trade-offs, and distribution of lifetime reproductive success.

## 2 Model and Methods

We begin with integral projection model (IPM) for soay sheep and then describe sampling life-histories to evaluate tradeoffs and the distribution of lifetime reproductive success. IPMs are structured population models that use linear (or linearized) regressions to describe the expected phenotypic trait trajectories (Coulson 2012; Ellner et al. 2016). The IPM consists of four functions that describe how traits such as body size (*z*) influence survival, reproduction, growth and offspring size.

For soay sheep survival, stage transitions and reproduction during a single time interval depend on age and stage (Coulson 2012). Based on Coulson (2012) and Kentie et al. (2020), we model soay sheep demography in discrete time with population projection matrices (PPMs) structured effectively by stage and add an maximum age of death. That is, we assume that at each age, the stage-structured matrices have same survival and fertility rates until the sheep eventually die off at a certain age. Based on empirical observations, the maximum age of death is set to 16. This enables us to construct a block matrix with stage-structured matrices for each age.

Note that the models are parameterised using female sheep data since the growth rate of soay sheep population is known to be female-dominant (Coulson et al. 2001) and there is no limiting effect of the number of males for female reproductive output. We use previously published data and IPM for soay sheep (Coulson 2012; Kentie et al. 2020). The generalized linear functions for survival and reproduction are as follows:

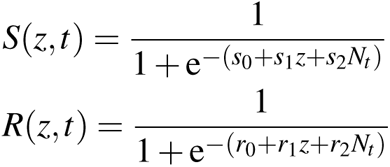

Here, *N_t_* is the population size at time *t*. *s*_0_, *s*_1_, *s*_2_ are the intercept, slope and coefficient for body size and density dependence and are our parameters of interest for varying survival. The same holds with *r*_0_, *r*_1_, *r*_2_ and recruitment function. For the parameter values, both survival and reproduction increase with body size *z* until it eventually saturates.

The growth *G* (*z*′|*z,t*) and parent-offspring (also called inheritance) *D* (*z*′|*z,t*)functions are described by Gaussian probability density functions. The growth function *G* (*z*′|*z,t*) is the probability that an individual with body mass *z* at time *t* will have a body mass *z*′ at time *t* + 1. The parent-offspring function *D* (*z*′ | *z, t*)is the probability that an individual with body mass *z* at time *t* will produce an offspring with body mass *z*′ at time *t* + 1.

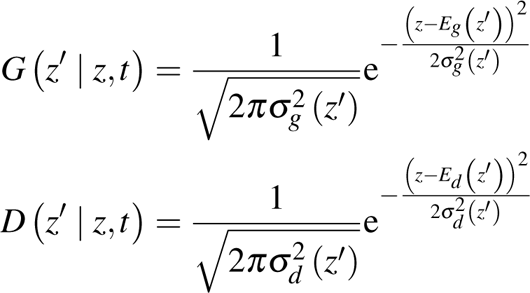

Here, *E_g_*(*z*′) is a linear function that predicts expected body mass and similarly *E_g_*(*z*′) predicts offspring’s expected body mass at *t* + 1 and are given by *E_g_* (*z*′) = *γ*_0_ + *γ*_1_*z* + *γ*_2_*N_t_* and *E* (*z*′) = *δ*_0_ + *δ*_1_*z* + *δ*_2_*N_t_*. 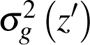 and 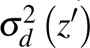 are also linear and are independent of population density. 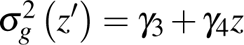 and 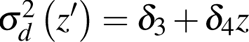. The table below summarizes the descriptions of our parameters of interest.

We define Growth Increment (GI) as average size in the next time minus size today:

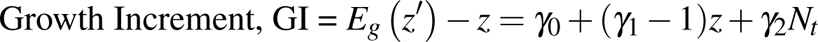

The intuition is that individuals reach an asymptotic size based on the stage (say y) at which growth increment (GI) becomes 0. We can then partition growth increment until stage y and beyond stage y for which individuals reach their asymptotic size. Thus the individuals yet to reach their asymptotic size will have a positive growth increment and is denoted by GIs. We refer to these individuals as juveniles in this manuscript. This is important to note because we then partition survival based on individuals who have not yet reached their asymptotic size and describe it as juvenile survival (as is described in the next section). The growth increment for individuals beyond their asymptotic size is denoted by GIb.

We use body mass distributed along 50 size classes as the trait since it is known to affect both survival and reproduction in Soay sheep, and is affected by population density. The IPM functions describe how body size z influences survival, reproduction, growth and offspring size by iterating the distribution of *z* at *t, n*(*z, t*) to *n*(*z*′, *t* + 1):

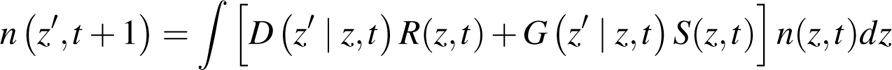

where population size is *N*(*t*) = ^∫^ *n*(*z, t*)*dz*. In matrix terms, the IPM can be written as:

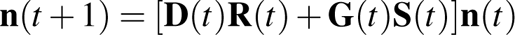

Using the IPM functions described above, we construct 50 × 50 stage-based matrices corresponding the each age for fertility and survival given by *F*(*z*′, *z*) = *D*(*z*′, *z*)*R*(*z*) and *P*(*z*′, *z*) = *G*(*z*′, *z*)*S*(*z*) (Steiner et al. 2014). We can construct a matrix **P_a_** which describes probability that an individual in stage *z* at age t is alive in stage *z*′ at t + 1 which is then used to calculate survivorship denoted by **L_a_**. Similarly, we get **F_a_** matrix which describes the number of stage *z*′ recruits produced by a female of size *z* at age *a*. The net reproductive rate *R*_0_ is calculated as the dominant eigenvalue of the matrix ∑*_a_* **F***_a_***L***_a_*. Note that the summation for ages *a* is until the age of death which is set to 16. In addition, we keep the matrices constant for all ages below the age of death which greatly simplifies our analysis. We can do this for different life-histories to understand the pattern of variation within the population. Below we describe how we generate many life-histories based on Coulson et al. (2006) and Kentie et al. (2020).

### 2.1 Average vital rate functions at SSD to evaluate tradeoffs

Coulson et al. (2006) introduced an approach called delifing or leave one out to estimate the contribution of an individual to realized changes in population size and stage-age distribution during a time interval. Using delifing, Kentie et al. (2020) estimated covariance across Soay sheep life-history parameters (such as *s*_0_, *s*_1_, … and so on) and examined how fluctuating population densities affect within-population variation in life-history strategies. The covariance structure for Soay sheep life history parameters is then used to span the range of possible strategies we expect sheep to follow given the phenotypic life-history trade-offs within and between demographic functions. We sample parameter sets of vital rates from a space of 10,000 life-history strategies (description of delifing in Coulson et al. (2006) and in Methods section of Kentie et al. (2020)).

A parameter set comprises of the 16 parameters: (*s*_0_, *s*_1_, *s*_2_, *r*_0_, *r*_1_, …), and are described in Table 1. For specific distribution (and values) for each of the parameters we refer to Kentie et al. (2020). We sample 200 such sets from the covariance matrix (using the delifing method). We calculate population growth rate given by *λ* as the dominant eigenvalue of the age-stage block matrix. The individuals are classified into 50 equal sized stage classes from 1 kg to 38 kg (based on their body size).

**Table 1:** The table provides a list of 16 parameters we use for the four IPM functions (Coulson 2012).

| Parameter | Description |
| --- | --- |
| $s_0$ | Survival: intercept |
| $s_1$ | Survival: slope body mass |
| $s_2$ | Survival: slope population size |
| $r_0$ | Recruitment intercept |
| $r_1$ | Recruitment: slope body mass |
| $r_2$ | Recruitment: slope population density |
| $\gamma_0$ | Development, mean: intercept |
| $\gamma_1$ | Development, mean: slope body mass |
| $\gamma_2$ | Development, mean: slope population size |
| $\gamma_3$ | Development, variance: intercept |
| $\gamma_4$ | Development, variance: slope body mass |
| $\delta_0$ | Inheritance, mean: intercept |
| $\delta_1$ | Inheritance, mean; slope body mass |
| $\delta_2$ | Inheritance, mean: slope population size |
| $\delta_3$ | Inheritance, variance: intercept |
| $\delta_4$ | Inheritance, variance: slope body mass |

We assume that the individuals reach a terminal age of 16 years and die. Thus the dimension of the block matrix is 800 × 800 because we have 50 size classes and 16 ages. For each parameter set, carrying capacity *K* is evaluated by finding the population size that results in *λ* converging to 1. Carrying capacity ratios (0, 0.5, 1) correspond to population size *N* = 0, *N* = *K/*2 and *N* = *K*, respectively. The results are robust with respect to sample and sample size. Figure 7 in the Appendix shows the range of equilibrium capacity for the 100 parameter sets.

To explore trade-offs, we calculated vital rate functions such as, juvenile survival (Ss), adult survival (Sb), Recruitment (R), and Growth Increment. We define Growth Increment (GI) as average size in the next time minus size today: GI = *E_g_ z*′ − *z* = *γ*_0_ + (*γ*_1_ − 1)*z* + *γ*_2_*N_t_*. We partition individuals before they reach asymptotic size based on the stage (say y) at which growth increment becomes 0 and take the average for growth increment until stage y. We denote the growth increment for these individuals as GIs. Similarly, Survival for those individuals (Ss) is evaluated by averaging upto the stage (y) and is called juvenile survival. Adult growth increment (GIb) and adult survival (Sb) are evaluated by averaging from (y+1) until the last stage. This way we generate a dataframe/matrix of average vital rates weighted by SSD for 200 parameter sets. Note that the function describing variance around the expectation 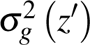 is independent of population density so we don’t consider it (we verified using calculations and the variance is small).

To understand trade-offs, we perform a PCA on average vital rate functions discussed above and the correlation plot at three densities. We evaluate the carrying capacity *K* for each parameter set described above and then calculate the average vital rate functions at the carrying capacity. This gives us a dataframe with 200 values for the 5 vital rate functions of interest. We can then repeat the exercise by calculating the average vital rate function at any ratio of carrying capacity. We predict the effects of density by focusing on population *N* = 0, *N* = *K/*2, *N* = *K*. We also explore the results from increasing the equilibrium capacity beyond *N* = *K*.

Since we are interested in examining the influence of population density on vital rate covariations, we note the derivative of *S*(*z, t*) and *R*(*z, t*) with respect to *N* is of the form:

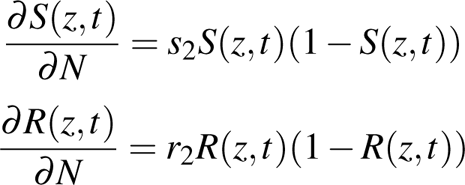

Since *S*(*z, t*) and *R*(*z, t*) are both logistic functions, they are always between 0 and 1. The values of *s*_2_ and *r*_2_ are negative and thus the derivative of *S*(*z, t*) and *R*(*z, t*) with respect to *N* will always be negative. We plot *S*(*z, t*) and *R*(*z, t*) for different values of *N* in the Appendix (Figure 8).

### 2.2 Lifetime Reproductive Success (LRS)

LRS measures the number of offspring an individual produces over its lifespan and individuals may produce 0, 1, 2… offspring, until they eventually die. The number of offspring produced during a time interval depend on the age and stage at the beginning of the time interval and is specified by probability distributions of producing 0, 1, 2, · · · offspring. The distribution is assumed to be Bernoulli since soay sheep produce either 0 or 1 offspring in a time interval and we ignore twinning (less than 1.9% of the offspring recruited are twins Simmonds and Coulson (2015)). Thus, following Tuljapurkar et al. (2020), we can compute the exact distribution of LRS for the age-stage-structured vital rates.

Tuljapurkar et al. (2020, 2021) developed an exact analysis to calculate the probability distribution of LRS for species described by age + stage models. The method assumes the empirical vital rates are known for a large cohort of individuals, for each st(age) of individuals life cycle. We use the method detailed in (Tuljapurkar et al. 2020) to compute LRS distribution of soay sheep individuals and examine demographic heterogeneities in the distribution.

The mean of LRS distribution is the net reproductive rate *R*_0_. We decompose *R*_0_ into probability of having no offspring and *R*_0_ conditional on making non-zero offspring, denoted by *γ* and is easily calculated as one minus the probability of reproductive failure (*β*_0_).

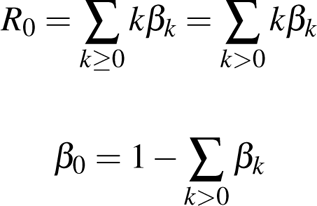

where *β_k_* is the probability of having *k* offspring. We are interested in *γ* = ∑*_k>_*_0_ *β_k_*, which is the mean LRS conditional on reproducing at least once.

## 3 Results

### 3.1 Trade-offs and Correlations at different densities

We examine trade-offs using average vital rate functions evaluated at SSD such as, juvenile survival (Ss), adult survival (Sb), Recruitment (R), Growth Increment for individuals before they reaching asymptotic size (GIs), and finally Growth Increment for individuals after reaching asymptotic size (GIb). We emphasize that juvenile survival (Ss) is calculated using the logistic function for Survival and is averaged until individuals reach their asymptotic size, corresponding to each parameter set (as described in the methods). Then we multiply by the SSD at that population density. The correlation plot at three densities (N=0, N=K/2, and K) is shown in the Appendix Figure 9.

We examined the trade-offs using PCA (Figure 1). 60-75% of the observed variation in vital rate estimates is explained by the first two principal components. The variance along the first principal component is explained by the trade-off between survival and growth until individuals reach their asymptotic size. However, at high densities the trade-off between survival and reproduction also becomes important as shown in Figure 1 and Table 2 (sign for Reproduction and juvenile survival are opposite). We validated our results using broken stick model and identified principal components whose eigenvalues surpass those predicted by the broken stick model (Figure 10 in Appendix). The variation captured by these components reflects significant underlying patterns in our data and is not a result of chance alone.

**Figure 1:**
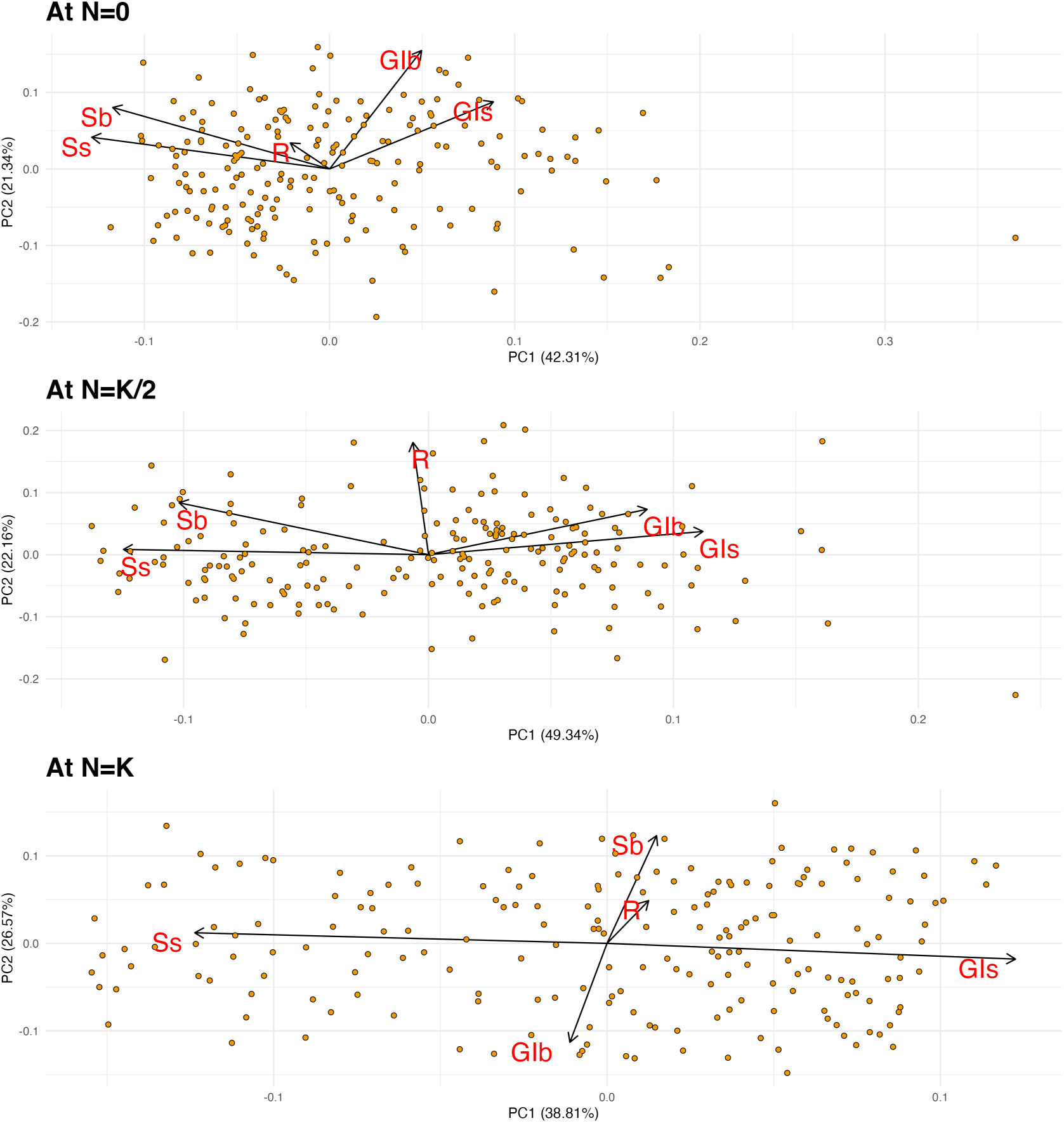
PCA results for average vital rate functions at SSD. Each panel corresponds to a density. Top most panel is for *N* = 0 case, followed by *N* = *K/*2, and *N* = *K*. Here Ss and Sb correspond to average survival (over 200 life-histories) for individuals before and after they reach their asymptotic size, respectively. Similarly, GIs and GIb is the average growth increment from one time to next for individuals before and after they reach their asymptotic size, and R is the average reproductive out-put.

**Table 2:** Table of PCA loadings for the five Principal Components.

| At N = 0 | PC1 | PC2 | PC3 | PC4 | PC5 |
| --- | --- | --- | --- | --- | --- |
| Ss | -0.635 | 0.205 | 0.16 | -0.04 | -0.727 |
| Sb | -0.579 | 0.397 | 0.052 | -0.298 | 0.645 |
| R | -0.105 | 0.168 | -0.973 | 0.083 | -0.08 |
| Gls | 0.437 | 0.433 | -0.018 | -0.756 | -0.223 |
| Glb | 0.245 | 0.765 | 0.154 | 0.576 | 0.004 |
| At N = K/2 | PC1 | PC2 | PC3 | PC4 | PC5 |
| Ss | -0.578 | 0.039 | -0.285 | -0.064 | 0.761 |
| Sb | -0.473 | 0.388 | -0.464 | 0.374 | -0.521 |
| R | -0.03 | 0.839 | 0.523 | -0.087 | 0.123 |
| Gls | 0.52 | 0.173 | -0.215 | 0.721 | 0.366 |
| Glb | 0.414 | 0.339 | -0.62 | -0.574 | 0.017 |
| At N = K | PC1 | PC2 | PC3 | PC4 | PC5 |
| Ss | -0.705 | 0.069 | -0.064 | -0.017 | -0.703 |
| Sb | 0.085 | 0.702 | -0.0005 | 0.706 | -0.033 |
| R | 0.071 | 0.278 | 0.905 | -0.29 | -0.119 |
| Gls | 0.698 | -0.102 | -0.12 | -0.015 | -0.698 |
| Glb | -0.063 | -0.644 | 0.403 | 0.645 | -0.052 |

Interestingly, we find strong negative correlation between Growth Increment (GIs) and Survival (Ss) for individuals before they reach asymptotic size is present at all densities. As we increase the density to equilibrium capacity (K), the strength of the negative correlation between growth increment and survival further increases (Figure 11 in Appendix). Thus, the trade-off between growth increment and survival is the largest at high population densities. In addition, a negative correlation between recruitment (R) and survival for juveniles (Ss) reveals itself only in the density dependent case in the bottom 2 panels of the figure 9 (not present in the density independent case in top panel). The figure 9 shows that the intensity of the association between juvenile survival and growth increases with density. This association has an increasing importance in shaping life history variation with increasing density and that at high densities there is a three-way trade-off between juvenile survival, growth increment, and recruitment.

What happens beyond equilibrium carrying capacity? We examine the demographic consequences when the population density increases beyond the equilibrium carrying capacity (Figure 12 in the Appendix). We find that now the trade-off between growth and survival weakens but the negative correlations between reproduction and survival increase. Although the trends do not remain consistently as density is increased beyond 1.4 times the equilibrium capacity.

### 3.2 Variation in *R*_0_ at different densities

Having identified the dynamic nature of trade-offs with population density, we wanted to examine density effects on demographic measures such as the net reproductive rate (*R*_0_) (same as mean of LRS) and the stable stage distribution. The *R*_0_ is calculated as the mean of the LRS distribution as well as through the age-stage block matrix (results are consistent). As discussed in the methods section, the model is age and stage-based in the sense that at age 16 we set survival to 0.

Figure 3 depicts the decrease in variation in *R*_0_ as a function of population size and we find both mean and variance of *R*_0_ decreases as population size increases. Since both *S*(*z, t*) and *R*(*z, t*) are monotonically decreasing in N, *R*_0_ is also a monotonically decreasing function of *N*. The key point here is the extent of variation in *R*_0_ for a density-independent (*N* = 0) and density-dependent case. The life-histories with large *R*_0_ at low-densities are highly impacted.

**Figure 3:**
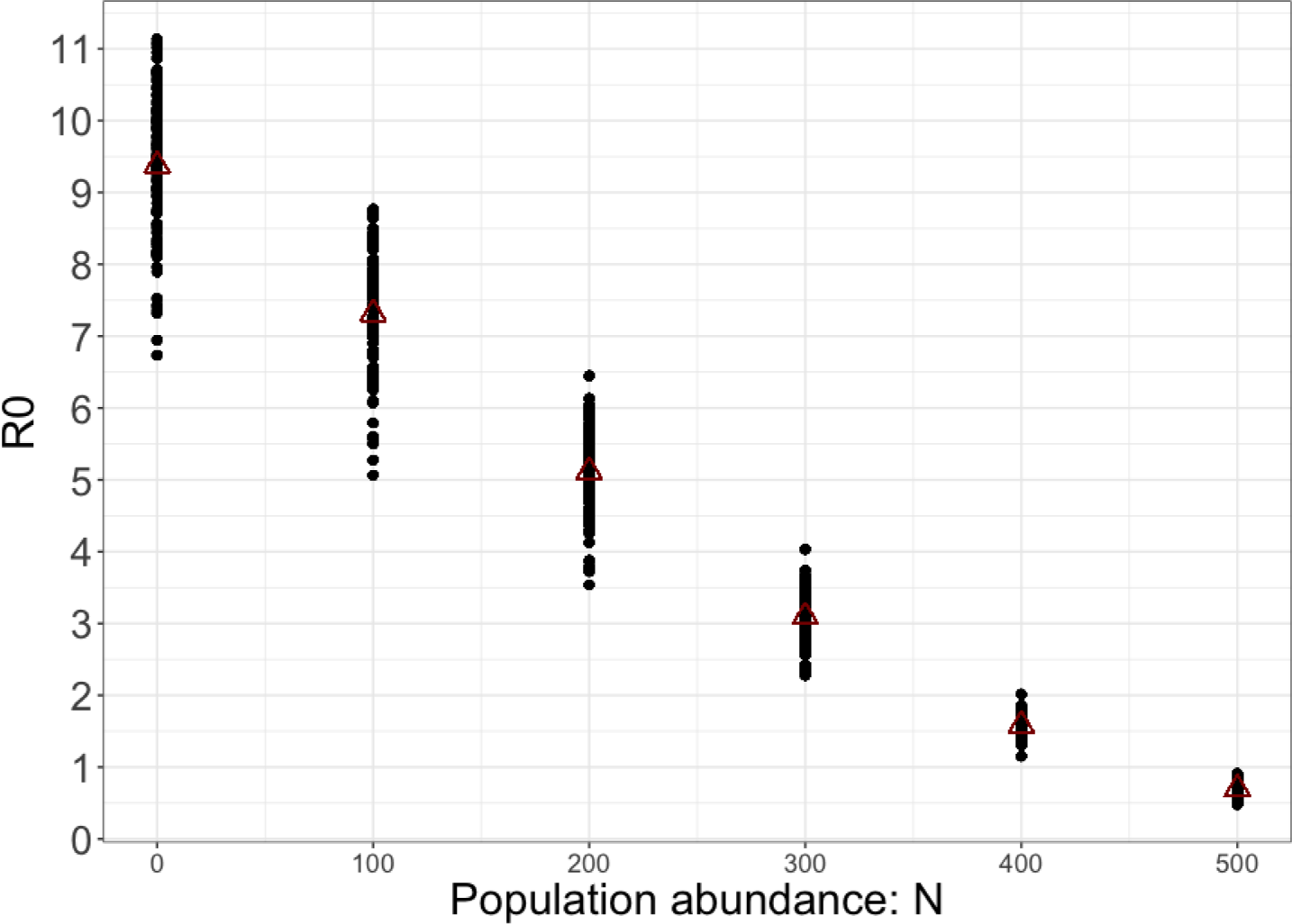
R_0_ as a function of population size

What factors contribute to decline in *R*_0_? We explore the question by understanding the variation in stable stage distribution (SSD) and mother’s size distribution at three population sizes (*N* = 0, *N* = *K/*2 and *N* = *K*). Since we are interested in sizes (stages), we examine the SSD for stage-based matrix population model. For each sample of parameter set, we estimate the equilibrium size *K* and calculate the stable stage (size) distribution (Figure 13 in Appendix). We take the average of SSD over 200 of our samples. Then we repeat by setting the population size at 0 and at *K/*2.

As population size increases from 0 to equilibrium population size *K*, the distribution/proportion of adults increases while the proportion of juveniles decreases, in contrast with density-independent case which has a higher proportion of juveniles (as shown in Figure 13 in the Appendix). We observe the same when we examine the distribution for mother’s size (scaled by SSD) as shown in Figure 4. We find the range of mother’s size to be less at higher densities thus contributing to lower *R*_0_. Thus, as density increases there is a concentration toward individuals of prime sizes (about 25kg) and contribution from individuals of sizes between 15-25kg is diminished. This is interesting because not only is there more competition and resource scarcity at high-density but there less individuals contributing to reproduction. Thus, the focus is to produce few extremely fit offspring and this aligns with r-K selection theory.

**Figure 4:**
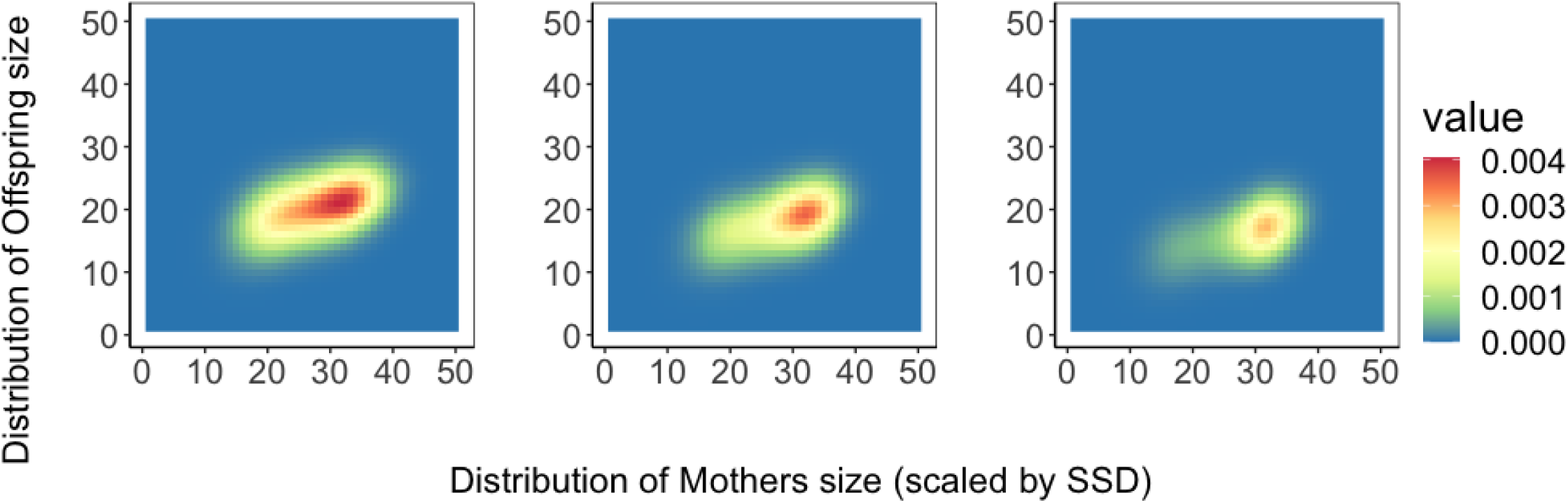
Plot for the distribution of Mothers size: on the x-axis we have mother’s size distribution scaled by SSD and on the y=axis we have offspring size. Each panel corresponds to a density. Left most panel is for *N* = 0 case, followed by *N* = *K/*2, and *N* = *K*.

Lastly, we examine the relationship between *R*_0_ at density-independence (by setting *N* = 0 to evaluate the matrices) and the equilibrium size *K* for the parameter sets. Figure 14 (in the Appendix) shows *K* and *R*_0_ are slightly negatively correlated as found in Kentie et al. (2020). Our results align with Pande et al. (2020, 2022) as they show mean population growth rate when rare (a measure of invasibility Chesson (2003)) is limited in its use as a metric for persistence.

### 3.3 Variation in vital rates

This section examines the relationship between average vital rate functions at SSD and mean lifetime reproductive success conditional on making at least one offspring (given by *γ*). We evaluate average juvenile survival (weighted by SSD) for our parameter set. Juvenile survival is calculated using the logistic function for Survival and is averaged until the individuals reach their asymptotic size (as described in the methods) for each of the parameter sets. Then we multiply by the SSD at that population size and regress the values of average juvenile survival at SSD against *γ* evaluated at the three population sizes.

In the three panels of Figure 5, we examine the relationship between average juvenile survival at SSD with the expected LRS conditional on making at least one offspring denoted by *gamma*. We find although increase in juvenile survival increases *R*_0_ and *γ*_0_ at low densities, near equilibrium size, *R*_0_ and *γ*_0_ are indifferent to changes in juvenile survival. We repeat as above by considering the relationship between average recruitment (at SSD) and *R*_0_ and *γ*_0_ and the results are similar to that for juvenile survival. We do not find any detectable relationship for adult survival and growth increment with *R*_0_ and *γ*_0_. Figure 5 show the constraints operating at high densities and equilibrium population sizes and perhaps how the mechanisms operating at different population sizes are different.

**Figure 5:**
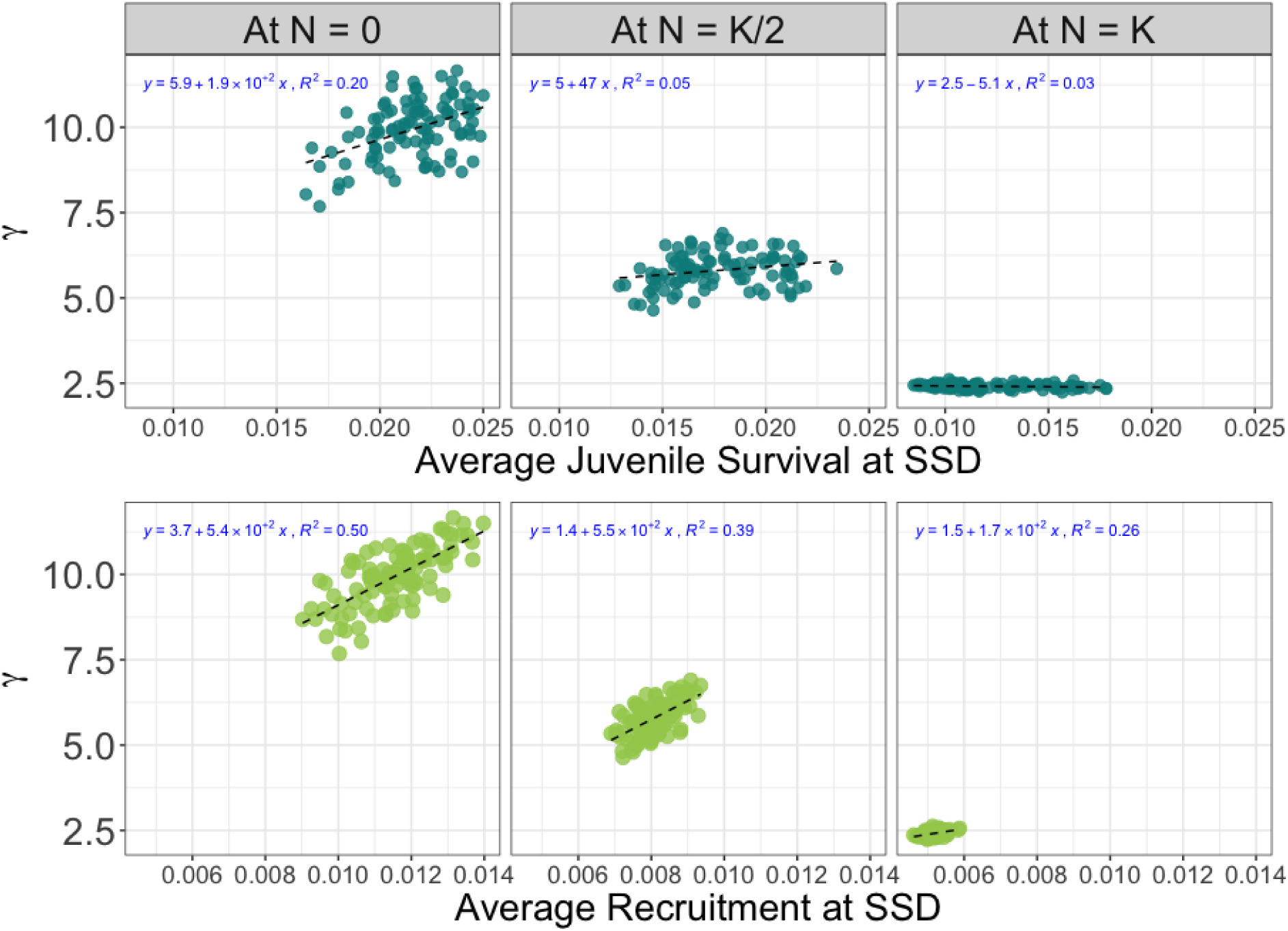
Trends for average vital rates at different population densities. On the y-axis is *γ*, that is the mean lifetime reproductive success conditional on making at least one offspring. The top three panels have average survival rates at SSD and the bottom three panels have average recruitment at SSD.

### 3.4 Variation in LRS trajectories at different densities

Now that we explored the effects of density on the mean LRS, we examine the entire distribution of LRS. We calculate the distribution of LRS (Tuljapurkar et al. 2020) for the 200 life-histories sampled from the covariance matrix. We hypothesized the variance of LRS at each age would be more constrained at higher population densities and indeed that is the case (Figure 15 in Appendix). The LRS trajectories show considerable variation at *N* = 0 which is not reflected at high densities since the size distribution is skewed toward individuals of large body sizes. Our results are consistent with empirical observations since we find high juvenile mortality at equilibrium compared to the density independent case.

To examine trade-offs from LRS distributions, we generated life-histories where survival parameters (*s*_0_, *s*_1_, *s*_2_) are sampled from the covariance matrix and the rest of the 13 parameters are fixed to their mean values. Similarly, we fixed all parameters to their mean valyes but recruitment parameters (*r*_0_, *r*_1_, *r*_2_). In the top panel of Figure 6 shows the average distribution of lifetime reproductive success and the confidence intervals for each offspring number. In the bottom panel of 6, we find at low population density, despite fixed survival small variation in fertility by chance alone can lead to higher or lower probability of having zero offspring. In the Appendix (Figure 15), we also calculate the distribution when all parameters are perturbed and our results remain consistent.

**Figure 6:**
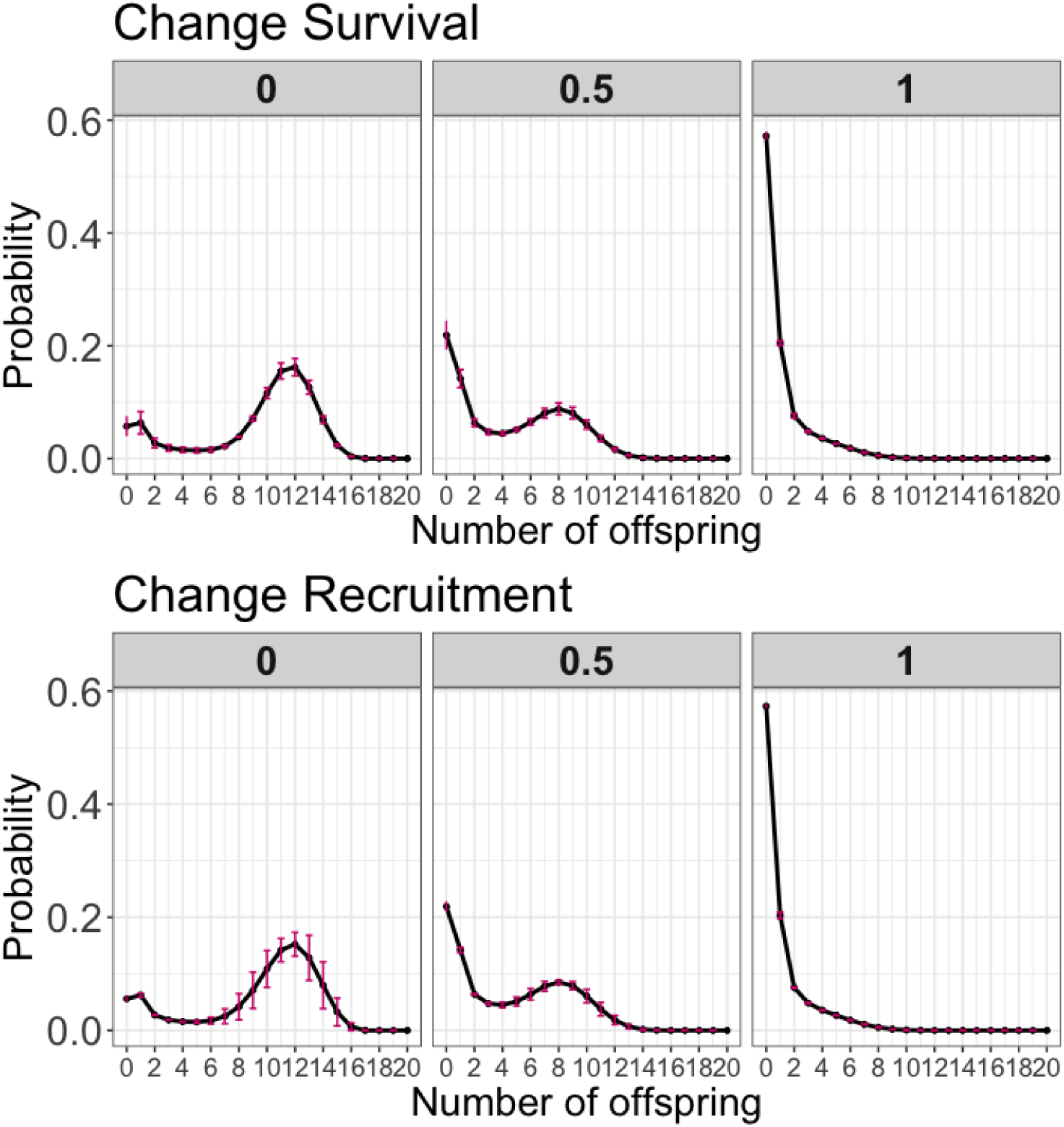
Plot for distribution of LRS. The top-three panels correspond to LRS distribution when only survival parameters are sampled from the covariance matrix and the rest of the parameters are fixed to their mean values. The bottom-three panels correspond to LRS distribution when only recruitment parameters are sampled and the rest of the parameters are fixed to their mean values. Each panel corresponds to a density. Left most panel is for *N* = 0 case, followed by *N* = *K/*2, and *N* = *K*.

Finally, we fixed all parameters but growth parameters. Our expectation was the additive effect of survival (only) and growth (only) would be greater than the effect of varying both survival and growth parameters together. We find slight evidence for such a trade-off for low offspring (0, 1 and 2) but further analysis is needed to uncover the trade-off (Figure 16 in Appendix).

## 4 Discussion

We examine the interplay between life-history trade-offs and density-dependence in Soay sheep. We use data from Kentie et al. (2020) to generate life-histories and found strong evidence for a trade-off between survival and growth increment for individuals before they reach their asymptotic size (Figure 9 and Figure 1 in Appendix). Interestingly, the nature of trade-offs is not static. The negative correlations increase as population density increases. In addition, at equilibrium carrying capacity a new trade-off arises between reproduction and juvenile survival. Thus, not all vital rates are equally impacted by density-dependent influences which gives rise to trade-offs. A demographic trade-off between growth and survival can be viewed as an axis of life-history variation where individuals can be placed along a slow-fast continuum (Stearns 1983; Oli 2004; Gaillard et al. 2016; Jiang et al. 2022; Van de Walle et al. 2023; Salguero-Gómez et al. 2016). Our findings are consistent with theory of life-history evolution (Stearns 1992) in that we find the evidence for trade-offs which prevent individuals from being proficient at both surviving and growing to large sizes.

Our results align with Kentie et al. (2020) as we find individuals with an average life history can not simultaneously maximise fitness for both high- and low-density environments. This is because demographic measures such as average survival, reproduction at SSD and distribution of lifetime reproductive success are highly constrained at high population densities (Figure 6). The level of variation and diversity at low population densities is not reflected at high population densities. Ozgul et al. (2009) found population density and maternal body size explain significant amounts of variation in temporal trends of mean body weight in Soay sheep. We find the contribution to reproduction for individuals at high population density comes from females of prime size of about 25kg (Figure 4). However, at low densities reproductive contribution is from females of both small and large sizes.

There is a concentration of individuals of prime size at high densities which is also revealed in our examination of stable stage distribution (Figure 13 in Appendix). Increase in density may be a double whammy for females of small sizes since there is more competition and resource scarcity and additionally there are less individuals contributing to reproduction. However, at high densities individuals of large body size may allocate in producing few extremely fit offspring at high population densities.

The results for probability of reproductive failure (*i.e.,* probability of making 0 offspring) also support empirical evidence within Soay sheep population in the wild. At low population densities, the juvenile mortality in Soay sheep is lower than the juvenile mortality at high population densities and this is reflected in the distribution of LRS in Figure 15. LRS is an important component of individual fitness and our research predicts how density dependent influences strongly shape the distribution of LRS and may potentially lead to density-dependent selection. Our research adds to the potential evidence for density-dependent effects on reproductive effort in natural populations, and aligns with r-K selection theory. A classic example is of Trinidadian guppies (*Poecilia reticulata*) that live in high population densities in the absence of predators. These guppies without predators compete more strongly for food, they mature later and exhibit lower reproductive effort in comparison to those living with predators at low population densities (Bassar et al. 2013). More recently, Travis et al. (2023) provide a comprehensive review of density-dependent selection and how it may promote contrasting patterns of trait means at different population densities.

There is empirical evidence for links between life-history trade-offs and density dependent selection from Drosophila cultures (Mueller and Ayala 1981; Mueller et al. 1991), insects (Gilbert and Manica 2010) and fisheries (Goodwin et al. 2006; Eikeset et al. 2016; Christie et al. 2018), among others.

Mueller and Ayala (1981) studied populations of *Drosophila melanogaster* kept at low population densities (r-populations for about 200 generations) and then placed them in crowded cultures (K-populations). They found after 25 generations the K-populations showed higher growth rate and productivity at high densities (relative to the controls), but lower growth rate at low densities and experimentally confirmed fitness trade-offs can arise from density-dependent selection. A study (Sæther et al. 2016) on great tits *Parus major* showed females laying the largest clutch sizes at small population sizes experienced the greatest density-dependent reductions in fitness at large population sizes, thus providing empirical support for r- and K-selection. At small population sizes, phenotypes with large growth rates are favored whereas phenotypes with high competitive skills are favored when populations are close to the carrying capacity K.

It is important to note that our results are based on life-histories generated from the covariance matrix in (Kentie et al. 2020) and don’t correspond to data from real individuals. We also do not take into account climatic variation and environmental stochasticity that have been found to affect population structure in Soay sheep (Ozgul et al. 2009). Lande et al. (2017) show (under assumptions for vertebrate populations) that the stochastic dynamics of population size can be accurately approximated by a univariate model governed by three key demographic parameters-the intrinsic rate of increase and carrying capacity in the average environment, and the environmental variance in population growth rate. In addition, while examining the distribution of LRS we have not considered the timing of reproduction which may be an important factor for density-dependent selection. Further exploration of trade-offs and density effects can account for timing of reproduction as well as age-dependence of demographic parameters, especially senescence for which there has been a compelling evidence for many ungulate populations (Loison et al. 1999; Gaillard and Lemaıtre 2019).

## 5 Appendix

**Figure 7:**
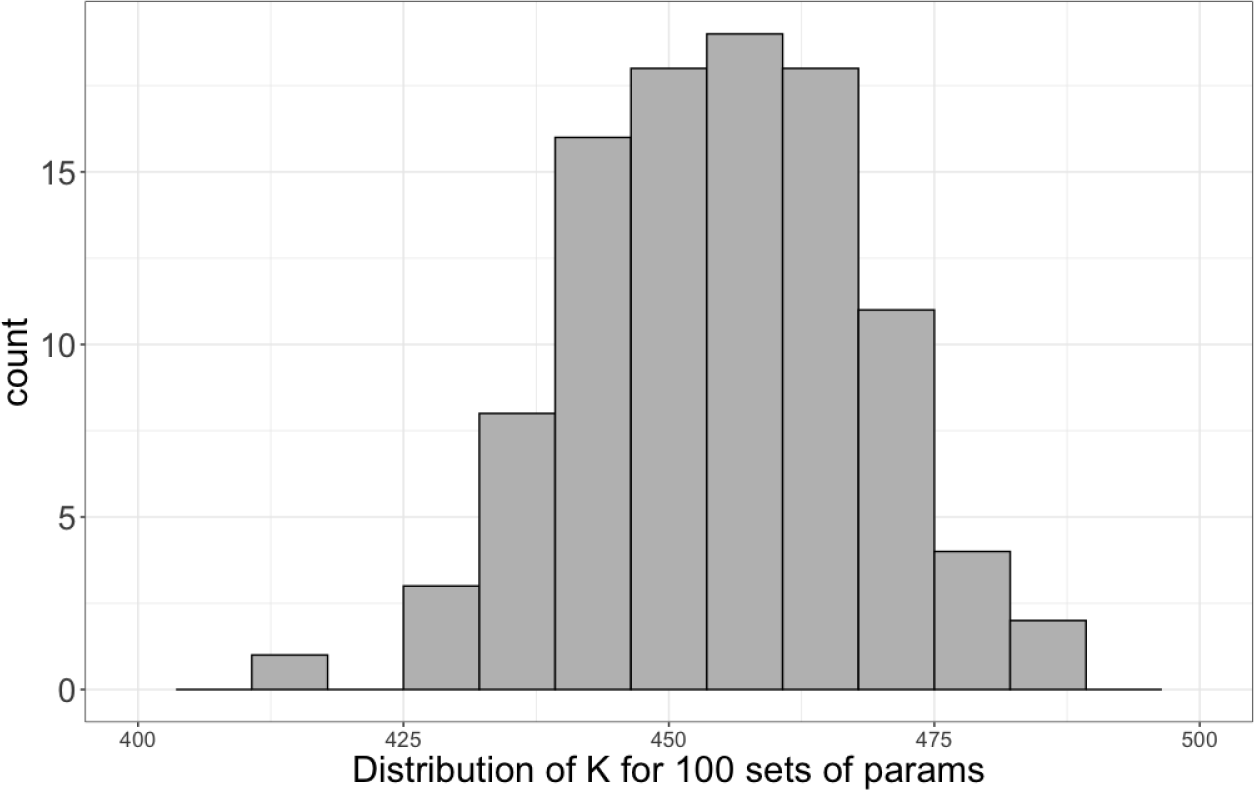
Equilibrium Distribution: mean 454

**Figure 8:**
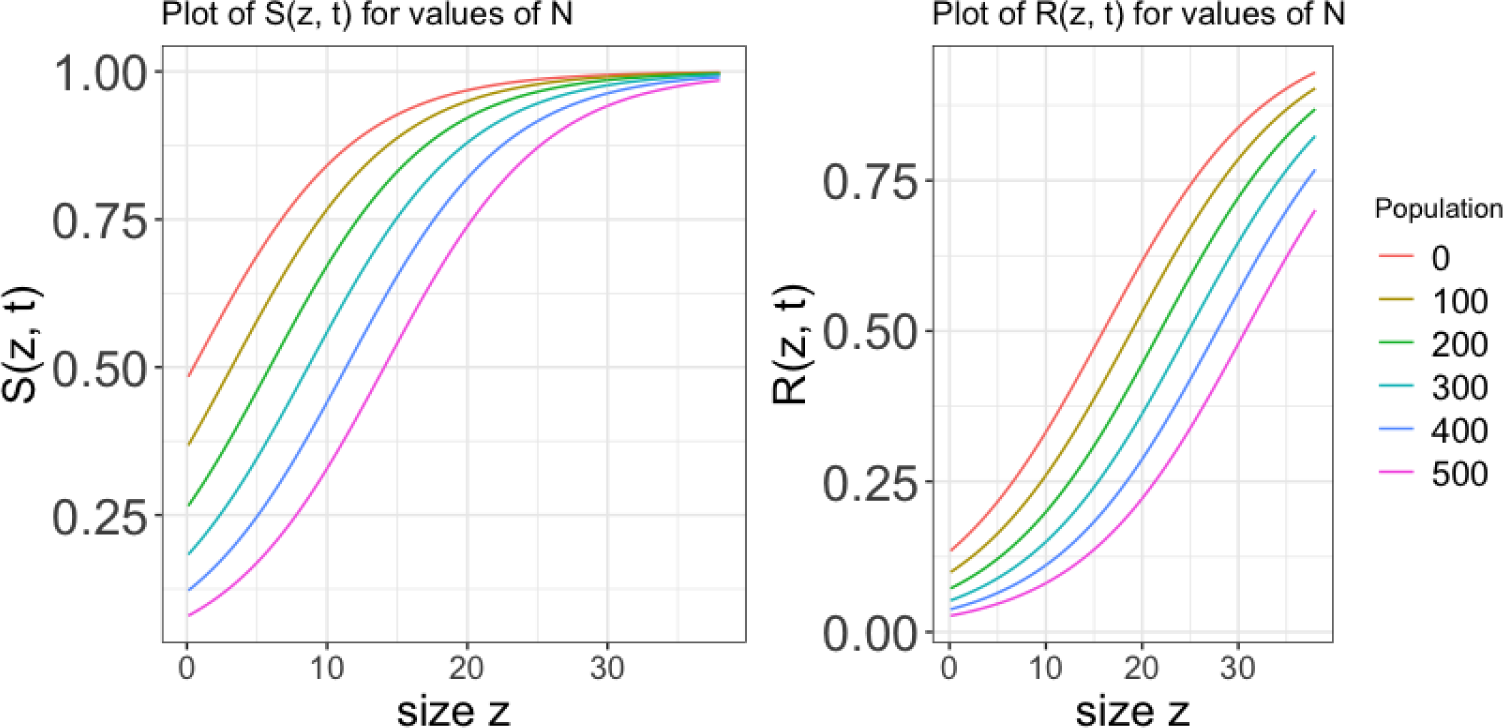
Plot of Survival and Recruitment functions for different values of Population, *N*.

**Figure 9:**
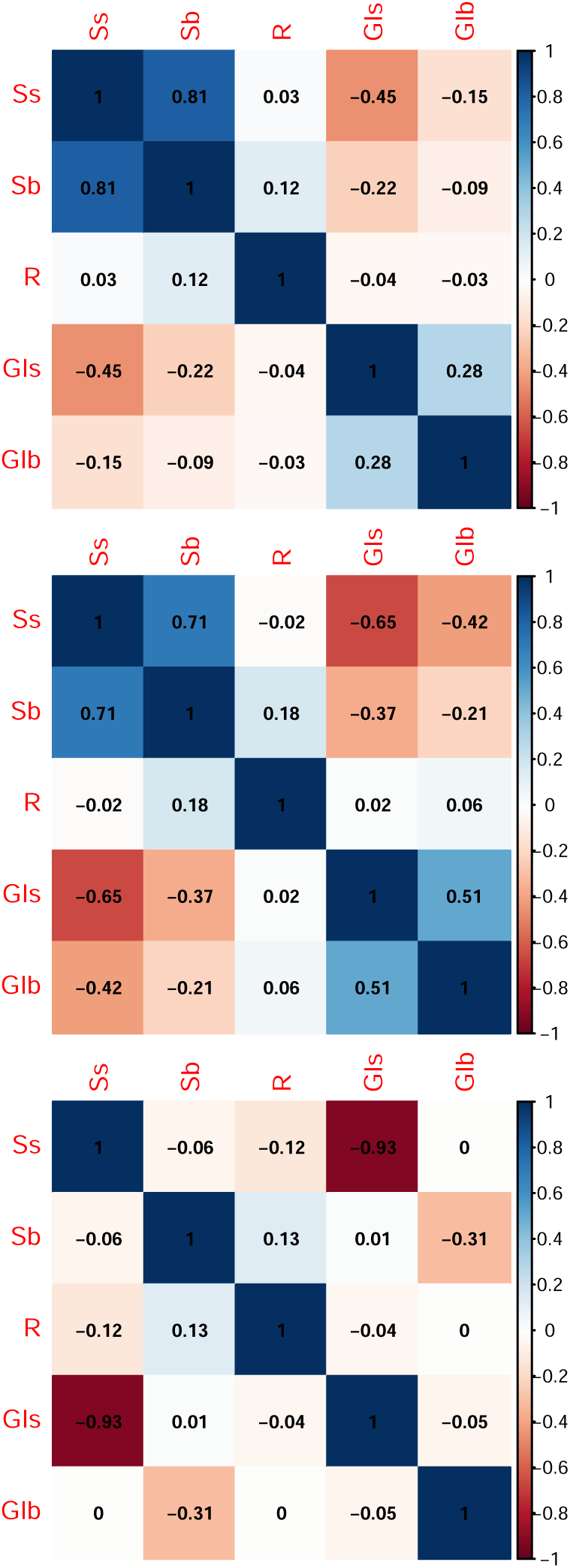
Correlation Matrices for the three population densities. The top most panel corresponds to *N* = 0, followed by *N* = *K/*2, and *N* = *K*. The colors red, blue, and white correspond to negative, positive and no correlation.

**Figure 10:**
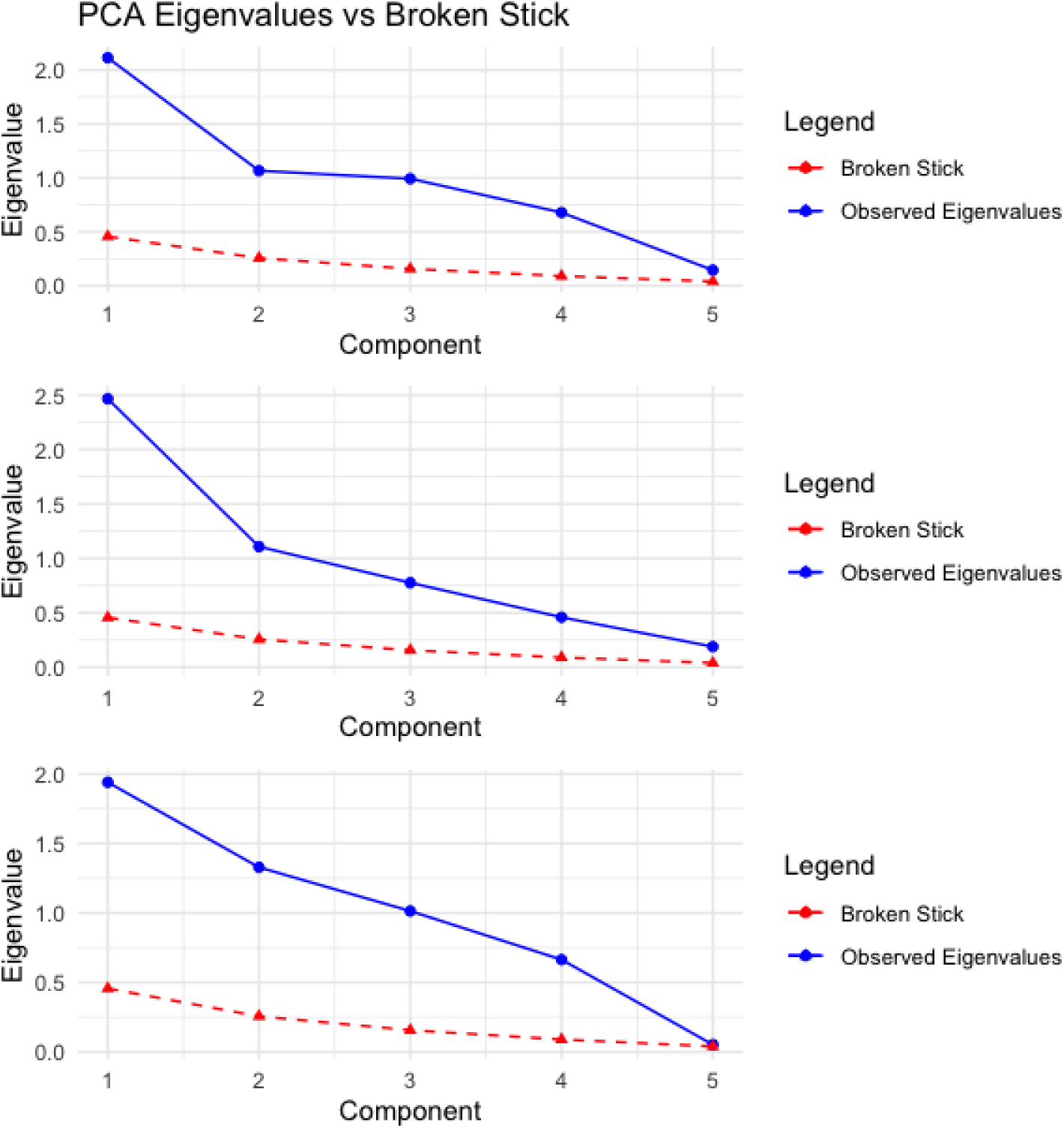
Eigenvalues from data and broken stick model. The top most panel corresponds to *N* = 0, followed by *N* = *K/*2, and *N* = *K*. Blue corresponds to observed eigenvalues from the data and red corresponds to eigenvalues from the broken stick model.

**Figure 11:**
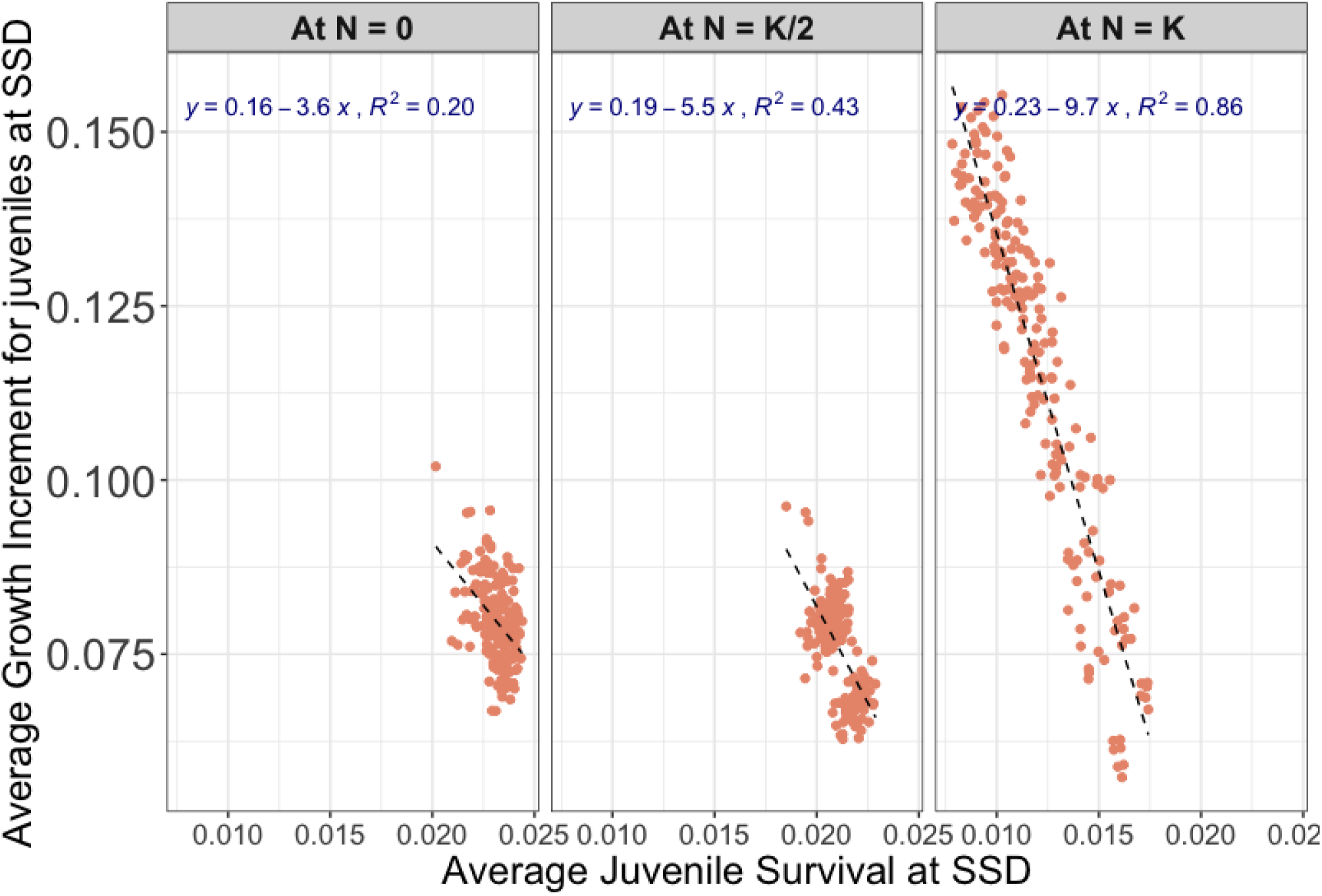
Plot for negative association between average growth increment for juveniles at SSD on the x-axis and average juvenile survival at SSD on the y-axis. The panels (left to right) corresponds to population density at *N* = 0, *N* = *K/*2, and *N* = *K*.

**Figure 12:**
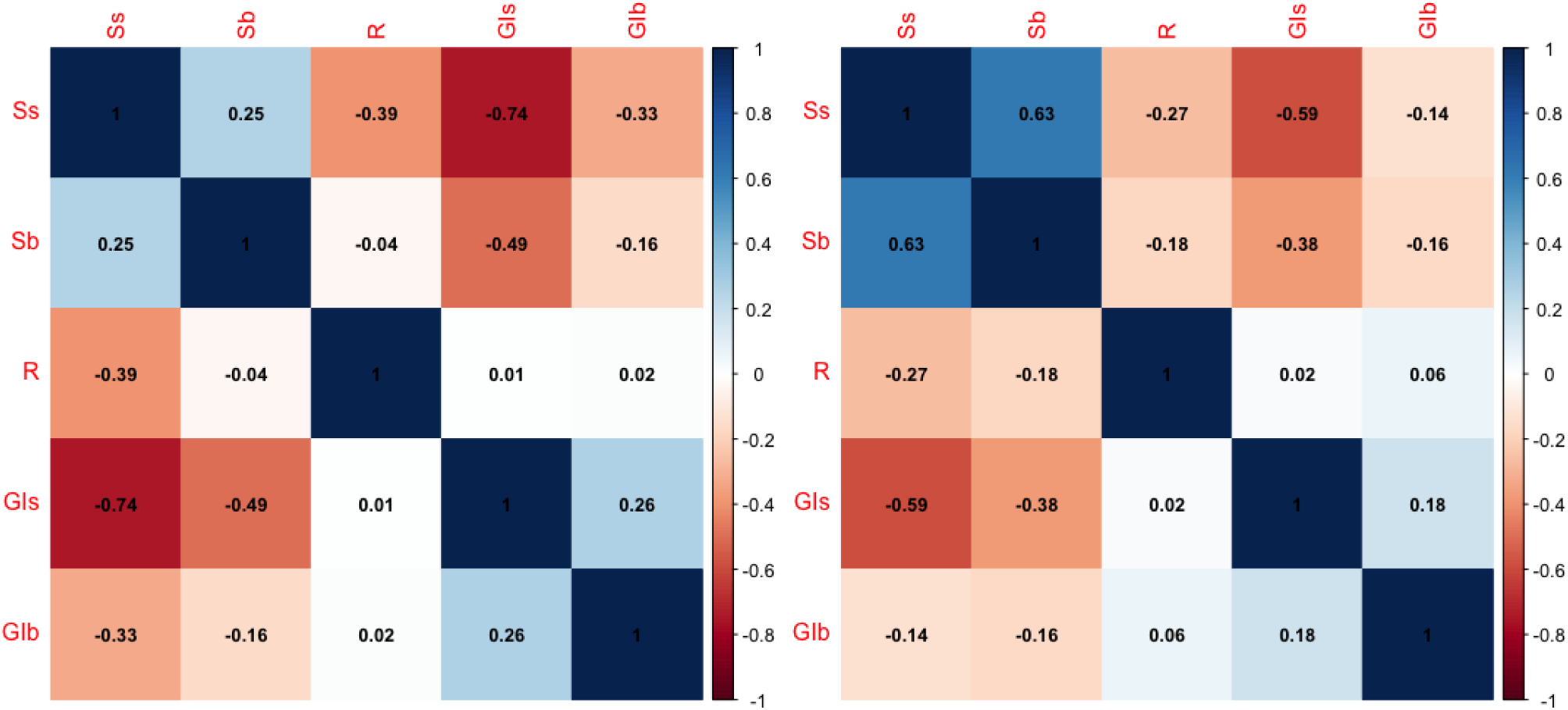
Correlation Matrices at *N* = 1.1*K* in the left panel and *N* = 1.3*K* in the right panel. The colors red, blue, and white correspond to negative, positive and no correlation.

**Figure 13:**
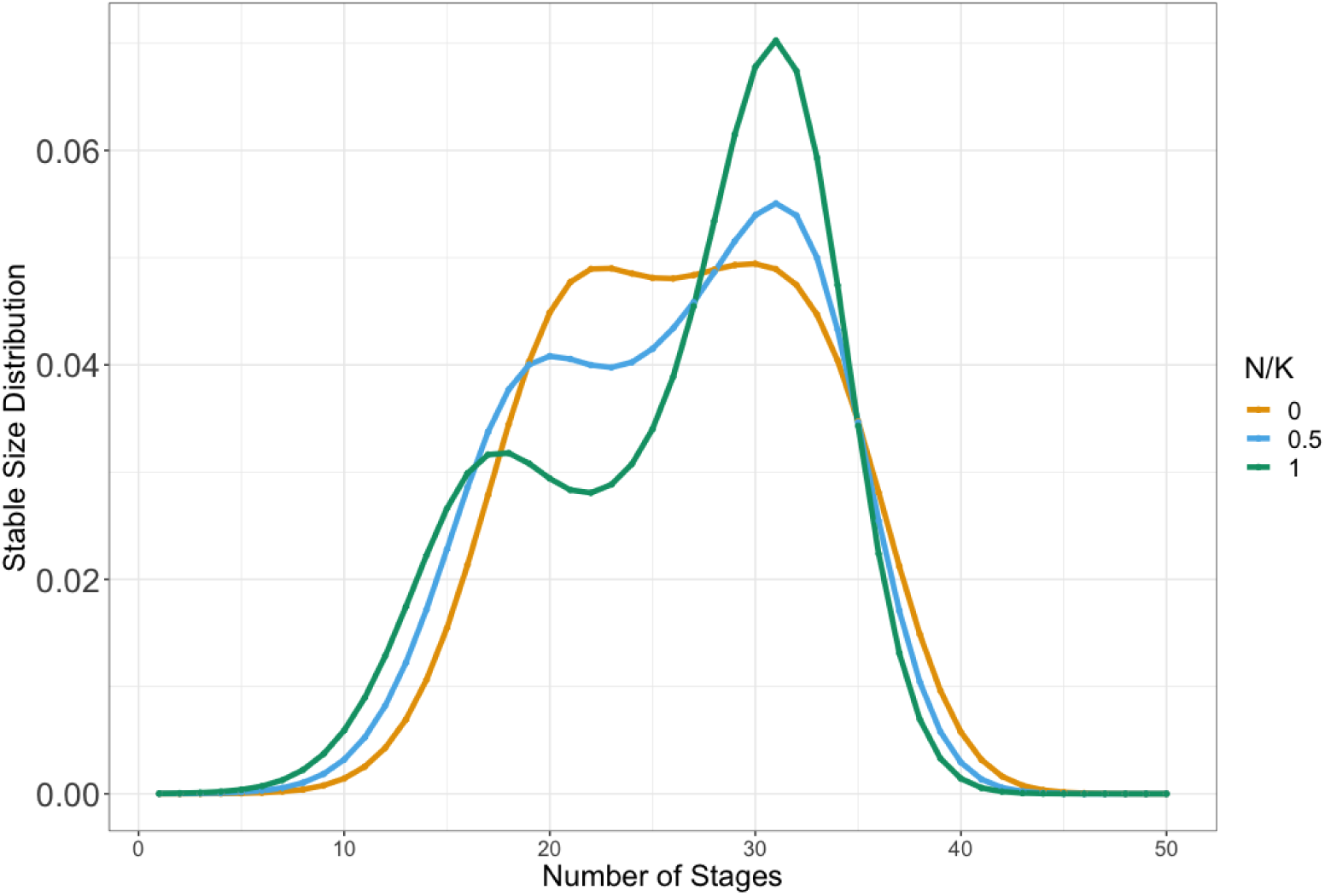
Stable Stage Distribution at different ratios of equilibrium population size. The legend *N/K* = 0, 0.5, and 1 correspond to *N* = 0, *N* = *K/*2, and *N* = *K*.

**Figure 14:**
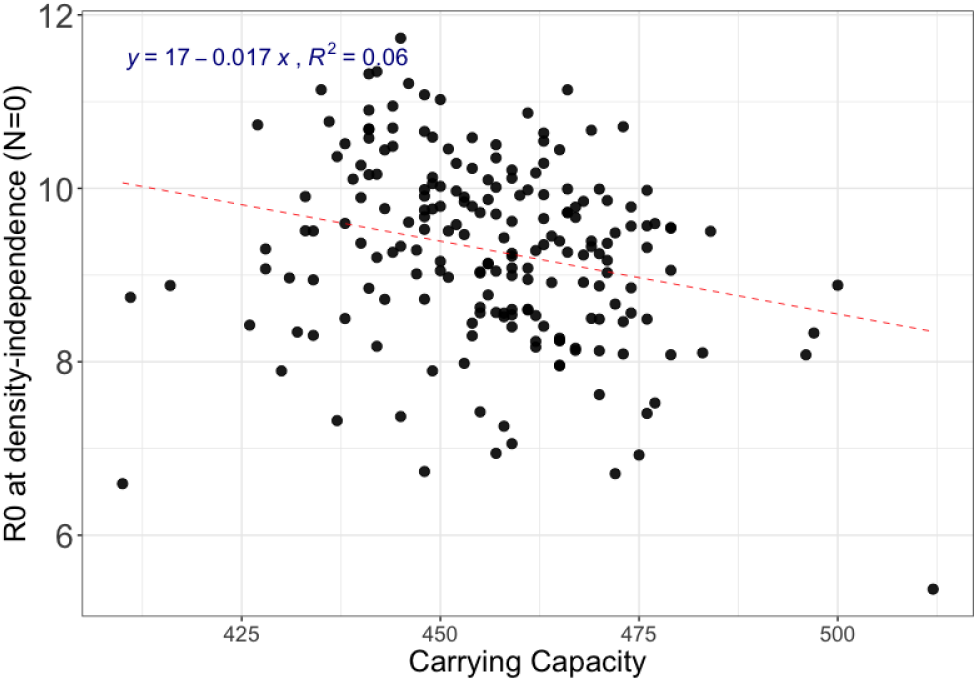
The plot shows negative correlation between carrying capacity, *K* on the x-axis and *R*_0_ at *N* = 0 on the y-axis.

**Figure 15:**
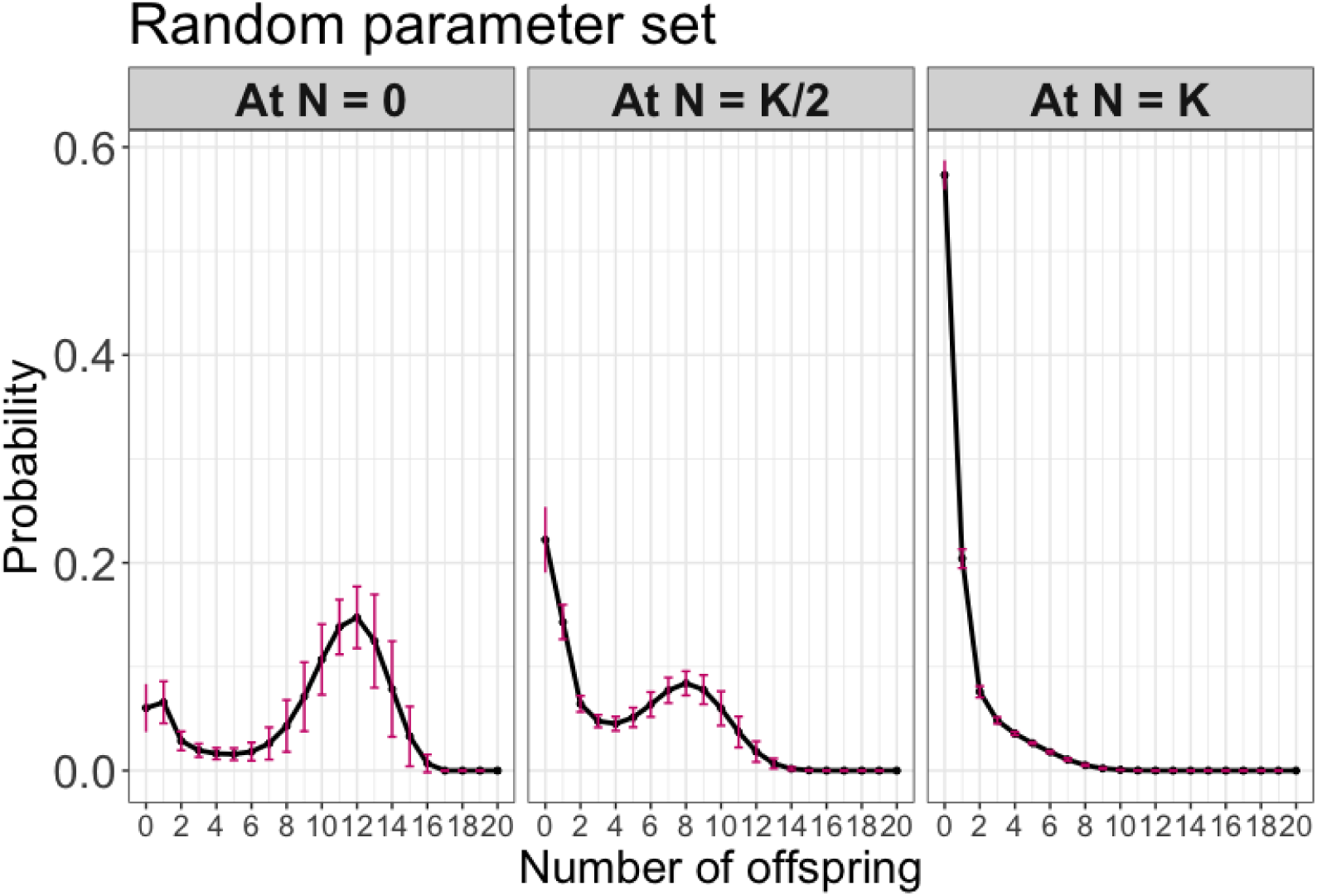
Distribution of LRS when all parameters are sampled from the covariance matrix at different population densities.

**Figure 16:**
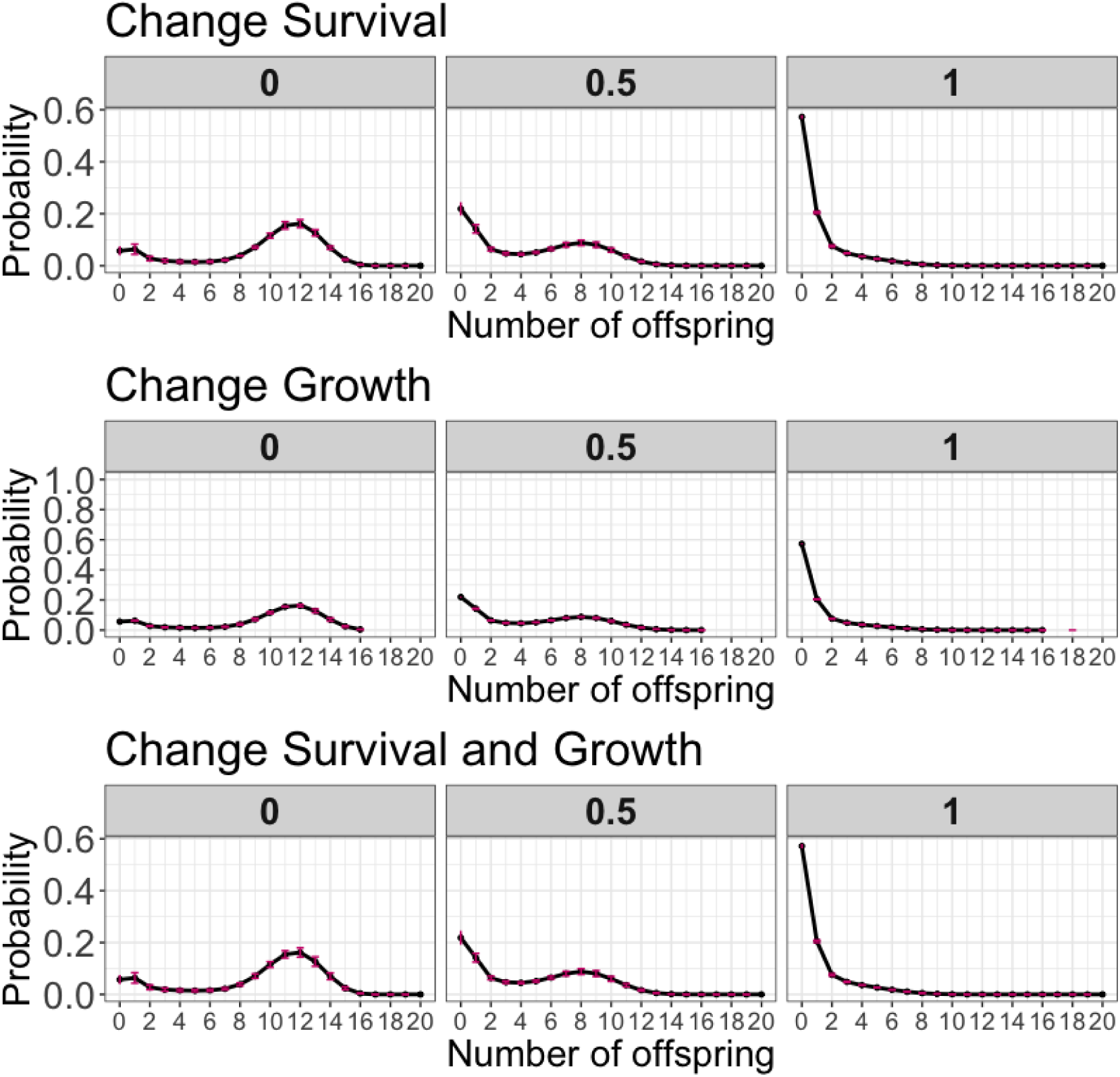
Distribution of LRS when survival and growth parameters are sampled from the covariance matrix at different population densities. The first and second panel corresponds to varying Survival and Growth parameters. The third panel corresponds to varying both survival and growth parameters simultaneously.

